# Stable sub-complexes observed *in situ* suggest a modular assembly pathway of the bacterial flagellar motor

**DOI:** 10.1101/369405

**Authors:** Mohammed Kaplan, Poorna Subramanian, Debnath Ghosal, Catherine M. Oikonomou, Sahand Pirbadian, Ruth Starwalt-Lee, Jeffrey A. Gralnick, Mohamed Y. El-Naggar, Grant J. Jensen

**Affiliations:** Division of Biology and Biological Engineering, California Institute of Technology, Pasadena, CA 91125.; Department of Physics and Astronomy, Biological Sciences, and Chemistry, University of Southern California, Los Angeles, CA 90089.; BioTechnology Institute, University of Minnesota – Twin Cities, St. Paul, Minnesota 55108.; Department of Plant and Microbial Biology, University of Minnesota – Twin Cities, St. Paul, Minnesota 55108.; Howard Hughes Medical Institute, California Institute of Technology, Pasadena, CA 91125.

## Abstract

The self-assembly of cellular macromolecular machines such as the bacterial flagellar motor requires the spatio-temporal synchronization of gene expression, protein localization and association of a dozen or more unique components. In *Salmonella* and *Escherichia coli*, a sequential, outward assembly mechanism has been proposed for the flagellar motor starting from the inner membrane, with each subsequent component stabilizing the last. Here, using electron cryo-tomography of intact *Legionella pneumophila*, *Pseudomonas aeruginosa* and *Shewanella oneidensis* cells, we observe stable outer-membrane-embedded sub-complexes of the flagellar motor. These sub-complexes consist of the periplasmic embellished P- and L-rings, in the absence of other flagellar components, and bend the membrane inward dramatically. Additionally, we also observe independent inner-membrane sub-complexes consisting of the C- and MS-rings and export apparatus. These results suggest an alternate model for flagellar motor assembly in which outer- and inner-membrane-associated sub-complexes form independently and subsequently join, enabling later steps of flagellar production to proceed.

## Introduction

In order to move efficiently in their low-Reynolds-number environment^1^, bacteria have evolved a complex membrane-embedded nanomachine known as the bacterial flagellum which exploits the flux of ions across the membrane to generate a mechanical torque to rotate a long filament^2–4^. In general, the bacterial flagellum consists of a cell-envelope-embedded motor, a hook and a filament. The motor consists of two parts: the rotor, which is composed of the basal body and the switch complex, and the stator (See ^5,6^ and references therein). In the canonical flagellar systems of *Salmonella enterica* and *Escherichia coli* the cell-envelope-spanning basal body consists of multiple rings: the MS- (membrane/supramembrane) ring (formed by the protein FliF), the P- (peptidoglycan) ring (FlgI), and the L- (lipopolysaccharide) ring (FlgH). The P- and L- rings form a bushing that allows the flagellum to rotate within the cell wall and outer membrane. These rings surround the rod structure (FliE, FlgB, FlgC and FlgF). The C- (cytoplasmic) ring consists of three proteins (FliG, FliM and FliN) and forms the switch complex which is responsible for switching the direction of flagellar rotation between clockwise and counterclockwise. In *E. coli* and *Salmonella*, the stator is formed by a complex of two proteins, MotA and MotB. While MotB interacts with the peptidoglycan through a peptidoglycan-binding domain^7–9^, MotA interacts with FliG to generate the torque required for flagellar rotation^8^. The extracellular parts of the bacterial flagellum comprise the hook, which acts as a universal joint, and the filament, which serves as a helical propeller. The motor also has a type III secretion system (T3SS) export apparatus consisting of six proteins (FliH, FliI, FliJ, FlhA, FlhB, FliP, FliQ and FliR) located at the inner membrane. See Fig. S1 for a schematic of these components.

High-resolution structures have been solved for many components of the flagellar motor by X-ray crystallography, NMR spectroscopy and cryo-EM single particle reconstruction, and structures of the purified *Salmonella* motor have been solved by electron microscopy^10,11^. Unfortunately, due to the large size of the complex and its integral position spanning the cell envelope, flagellar motors lose components when purified. Recently, the advent of electron cryo-tomography (ECT)^12,13^ allowed our group and others to reveal the *in situ* structures of various bacterial flagellar motors within intact cells at macromolecular (~ 4 nanometer) resolution. These studies showed the diversity of flagellar motors in different species adapted to unique external environments^14–18^. For instance, some species elaborate their P- and L-rings with additional periplasmic components, including the T-ring (MotX and MotY) and H-ring (FlgO, FlgP and FlgT)^19,20^.

The bacterial flagellum offers a striking example of the self-assembly process of supramolecular complexes in the cell, and is of interest to disciplines ranging from evolutionary biology to nanotechnology^21^. Assembly requires huge amounts of cell energy^22,23^. Our current understanding of flagellar assembly comes from studies of two enteric species of Gram-negative bacteria, *E. coli* and *Salmonella*, which suggest an inside-to-outside sequential assembly process starting from the export apparatus (FlhA) and MS-ring (FliF) followed by the C-ring proteins and the export apparatus, the rod, the P- and L-rings, the hook and finally the filament^22,24–26^. The process is thought to be a cooperative one whereby the addition of each new component stabilizes its antecedent^26^. The proteins forming the rod, the hook and the filament are secreted through the T3SS export apparatus^22^. The P-, L-, T- and H-ring proteins, however, are secreted to the periplasm through the conventional Sec pathway^22,27–29^. Interestingly, the P- and L-ring proteins in *S. typhimurium* were found to exist in a stable state in the periplasm in the absence of the inner membrane-associated complex^28^. Despite the stability and independent export of their components, it is thought that P- and L-rings form only once the assembling rod extends through the periplasm^22^.

By imaging intact cells of three non-enteric Gram-negative bacterial species, we describe here that the P/L-ring sub-complexes (with associated rings) and the inner membrane complex (constituting the C-ring, the MS-ring and the T3SS export apparatus) form stable independent sub-complexes, suggesting an alternative assembly model in which modules form independently and associate into a functional structure.

## Results & Discussion

We recently reported the structure of the intact flagellar motor in three non-enteric Gammaproteobacteria species, *Legionella pneumophila*, *Pseudomonas aeruginosa* and *Shewanella oneidensis* MR-1 by ECT^30^. *L. pneumophila* and *P. aeruginosa* are human pathogens that cause serious pulmonary infections in which the flagellum is a key virulence factor^31,32^. *S. oneidensis* is a model system for studying extracellular respiration and is known for its production of multiheme cytochrome electron conduits and outer-membrane appendages^33^. All three species utilize a single, polar flagellum. We found that the flagellar motors of all three species contain elaborated P- and L-rings: *L. pneumophila* and *P. aeruginosa* have an extra periplasmic ring surrounding the P-ring, and *S. oneidensis* has both T-and H-rings surrounding its P- and L-rings, respectively^30^.

In addition to fully-assembled flagella, in tomograms of all three species we observed isolated outer membrane complexes similar to the periplasmic P- and L-rings and their associated rings (henceforth these complexes of the P-, L-rings and associated rings are referred to as PL sub-complexes) (Fig. 1). By performing sub-tomogram averaging of these sub-complexes to enhance the signal-to-noise ratio we confirmed that these were PL sub-complexes and that they lacked other flagellar components (Fig. 1 C, D, H, I, M and N; compare to averages of fully-assembled motors in 1 E, J and O). Compared to fully-assembled motors, PL sub-complexes had two striking features. First, they sharply curved the outer membrane inward into an inverted omega shape. The membrane remained continuous, however, and no pore was visible. Second, in *L. pneumophila* and *P. aeruginosa* two protein densities were seen extending downwards from the center of the sub-complexes (Figs. 1 C, D, H and I, purple densities). These densities were less clear in the case of *S*. *oneidensis* (Fig. 1 M). We speculate that these densities may play a role in docking the PL sub-complex to more proximal components. Previous ECT studies of the flagellar motor in other species may not have observed PL sub-complexes because they lack the ornamentation of the T- and/or H-rings, which enhance visibility.

**Figure 1:**
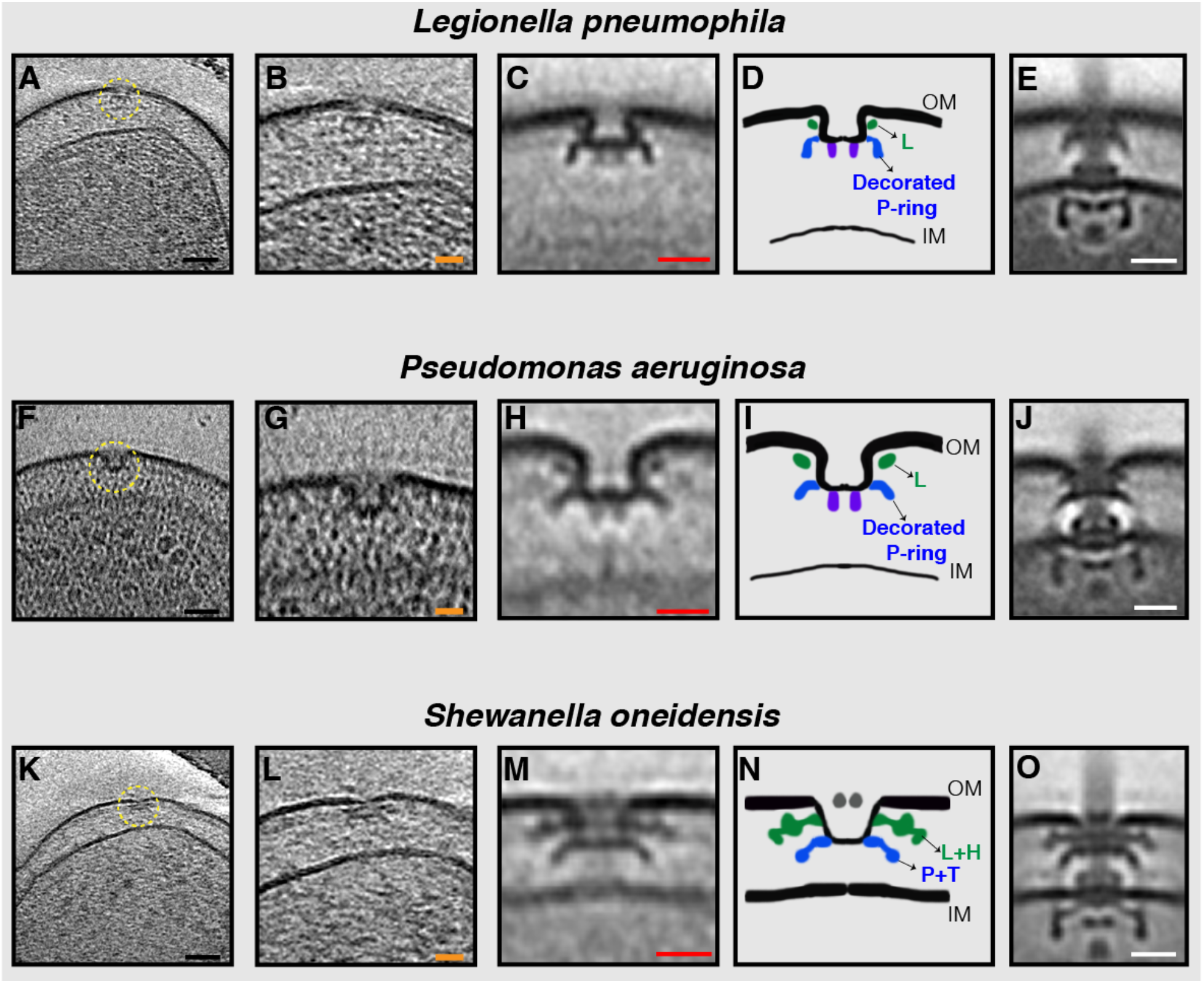
Stable PL sub-complexes in three bacterial species imaged by ECT. **(A, F, K)** slices through electron cryo-tomograms of *L. pneumophila*, *P. aeruginosa*, and *S. oneidensis* cells, respectively, highlighting a PL sub-complex in the outer membrane (dashed yellow circle). **B, G, L)** Enlarged views of the complexes. **C, H, M)** Sub-tomogram averages of PL sub-complexes from each species. **D, I, N)** Schematic representations of the sub-tomogram averages with different rings colored and labeled. **E, J, O)** Sub-tomogram averages of fully-assembled flagella from each species for comparison. Scale bars: (black) 50 nm, (orange) 25 nm, (red and white) 20 nm.

In many *L*. *pneumophila* cells, we also observed an inner membrane complex constituting the C- and MS-rings and the T3SS export apparatus (referred to henceforth as the inner-membrane (IM) sub-complex) in the vicinity of the PL sub-complexes (Fig. 2 A-L and Movies S1 and S2). The lateral distance between the PL sub-complex and the IM sub-complex ranged from 60 nm to 5 nm (Fig. 2 D-L and Table S1). We also observed that the distance between the inner and outer membranes varied and that this variation correlated with the lateral distance between the sub-complexes; the more closely aligned the two sub-complexes were, the closer the inter-membrane distance was to the distance observed in fully-assembled flagella (35 nm) (Table S1 and Fig. S2). We never observed an IM sub-complex without a PL sub-complex in its vicinity. In 11 *L. pneumophila* cells, we found fully assembled motors lacking the hook and filament, but no motors with only the hook (and not the filament) were observed. In one tomogram of a lysed *L. pneumophila* cell, we found an example of a flagellar sub-complex where only the rod, the outer membrane complex, the hook and the filament were present without the C-ring and export apparatus (Fig. 2 M and N). Since secretion of the rod, hook and filament proteins into the periplasm requires the T3SS export apparatus, this must represent an intermediate stage of disassembly. This pattern is similar to what was previously seen in *Caulobacter crescentus* where the disassembly process is initiated by digestion of the C terminus of FliF, leaving the rod, the hook and the filament as a stable sub-complex that is ejected into the medium^34–37^. In addition, the FliG and FliM components of the C-ring (which is connected to the export apparatus) were also shown to undergo proteolysis during the disassembly process in *C. crescentus*^34,38^.

**Figure 2:**
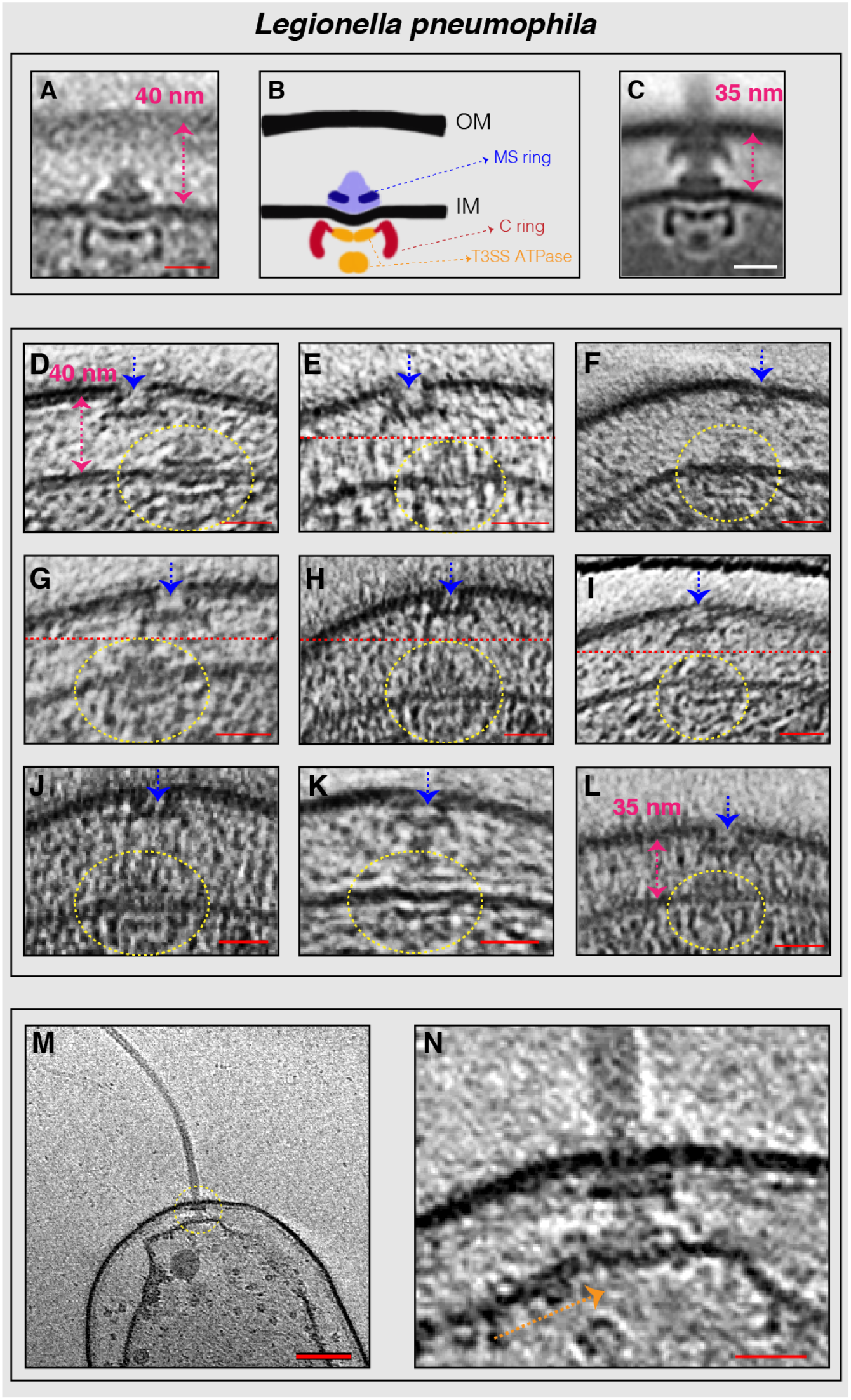
Flagellar sub-complexes in *L. pneumophila*. **(A)** Sub-tomogram average of the IM sub-complex constituting the C- ring, MS-ring and export apparatus. **B)** Schematic representation of the sub-tomogram average shown in (A) highlighting the different parts of the complex. **C)** Sub-tomogram average of the motor of fully-assembled flagella highlighting the distance between the inner and outer membranes. **D-L)** Slices through electron cryo-tomograms showing neighboring PL and IM sub-complexes. Dashed-yellow circles highlight the IM sub-complex while dashed-blue arrows highlight the PL sub-complex. Dashed-pink arrows highlight the distance between the inner and outer membranes. Dashed-red lines mark the border between two images used to make a composite image when the PL and IM sub-complexes were found at different Z-levels in the tomogram. **M)** Central slice through an electron cryo-tomogram of a lysed cell. The dashed-yellow circle highlights the flagellar motor. **N)** Enlarged view of the same slice shown in M. The absence of the C-ring and the export apparatus is highlighted by the dashed-orange arrow. Scale bars: (A, C) 20 nm, (D-L, N) 25 nm, (M) 100 nm.

In tomograms of *P. aeruginosa* cells, we found (next to fully-assembled flagella and PL sub-complexes) examples of fully-assembled motors both without (5 cases) and with (3 cases) the hook (Fig. 3). The low number of particles in this state suggests a fast transition from the fully-assembled motor to the fully-assembled flagellum. Similarly, in four *S. oneidensis* cells we saw fully-assembled motors lacking the hook, next to fully-assembled flagella and PL sub-complexes (Fig. 4 A-E).

**Figure 3:**
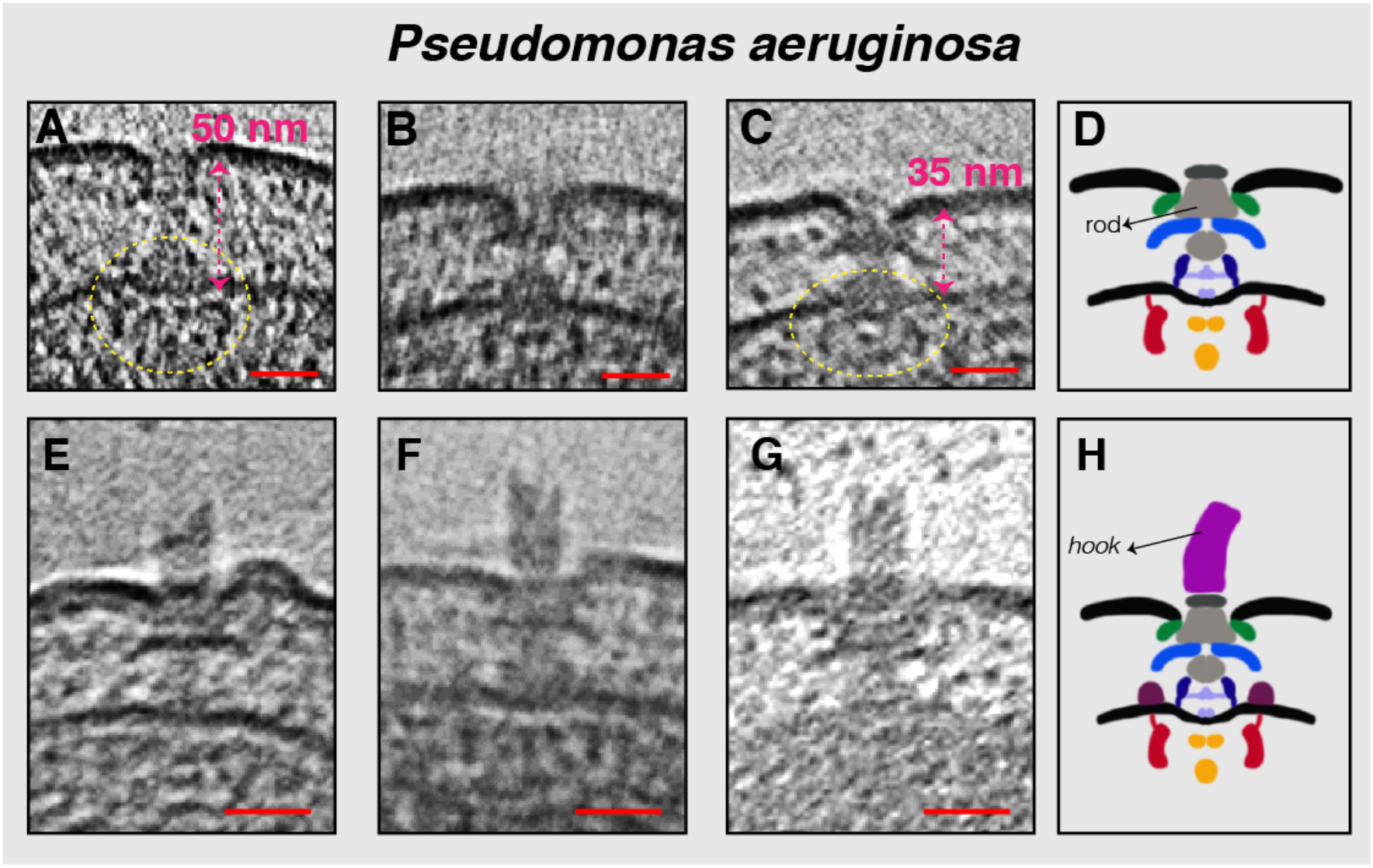
Flagellar sub-complexes in *P. aeruginosa*. **(A-C)** Slices through electron cryo-tomograms showing fully-assembled motors without the hook and filament. The distance between the inner and outer membranes is highlighted by the dashed-pink arrows. The dashed-yellow circles indicate the IM sub-complex. **D)** Schematic representation of the *P. aeruginosa* motors lacking the hook and filament shown in (A-C). **E-G)** Slices through electron cryo-tomograms showing fully-assembled motors with the hook and lacking the filament. **H)** Schematic representation of the motors with the hook shown in (E-G). Scale bars are 25 nm.

**Figure 4:**
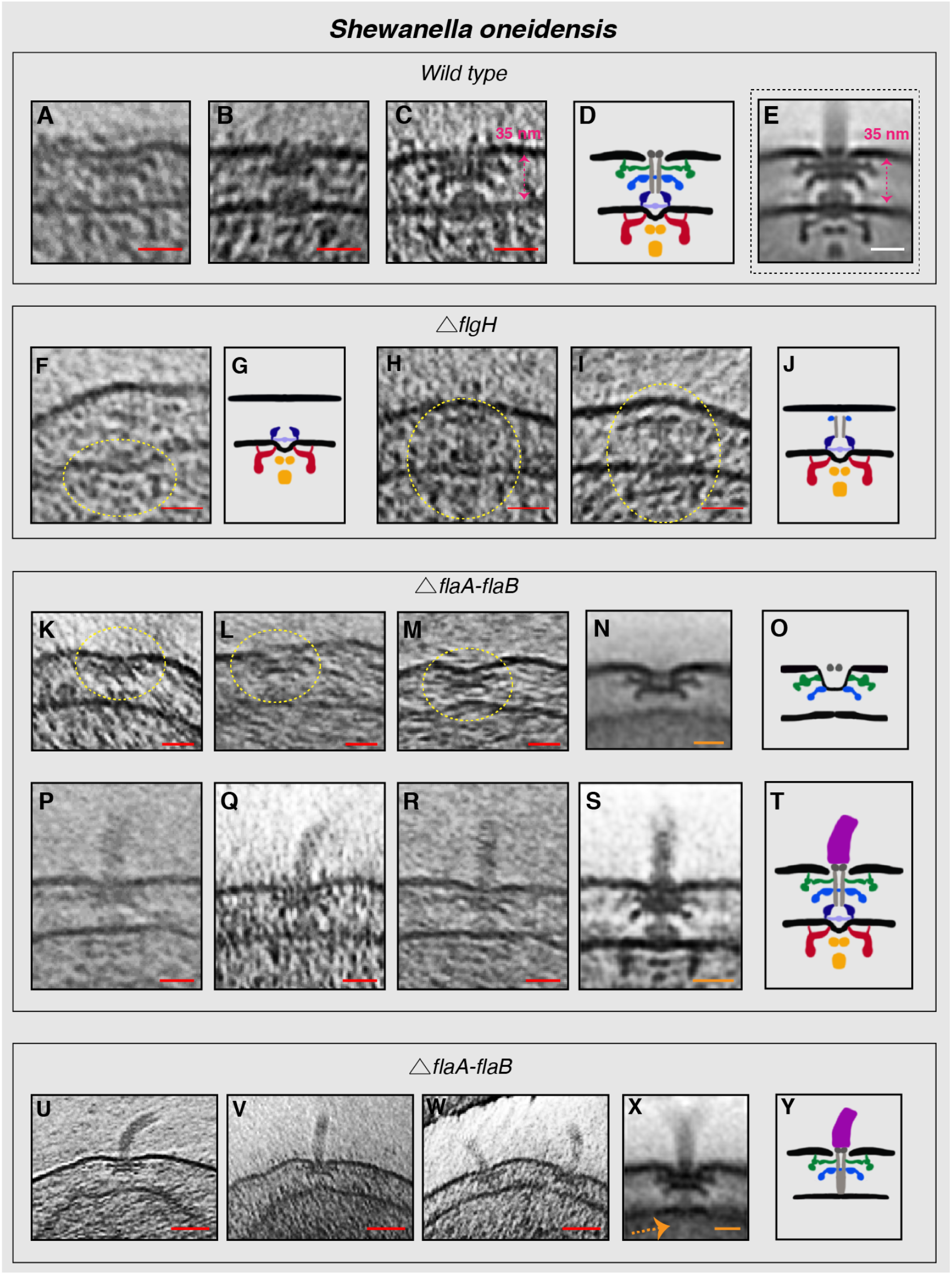
Flagellar sub-complexes in *S. oneidensis* wild type and mutant cells. **(A-C)** Slices through electron cryo-tomograms of wild type cells showing fully-assembled motors without the hook and filament. The dashed-pink arrow highlights the distance between the inner and outer membranes. **D)** Schematic representation of the motors lacking the hook and filament shown in (A-C). **E)** Sub-tomogram average of the motor of fully-assembled flagella with the distance between the inner and outer membranes highlighted by the dashed-pink arrow. **F)** Slice through an electron cryo-tomogram of a *∆flgH* cell showing an IM sub-complex, indicated by the dashed-yellow circle. **G)** Schematic representation of the IM sub-complex shown in F. **H & I)** Slices through electron cryo-tomograms of *∆flgH* cells showing the IM sub-complex with the rod and the P-ring, indicated by the dashed-yellow circles. **J)** Schematic representation of the structures shown in H and I. **K-M)** Slices through electron cryo-tomograms of *∆flaA/B* cells highlighting PL sub-complexes (dashed-yellow circles). **N)** Sub-tomogram average of the PL sub-complexes in *∆flaA/B* cells. **O)** Schematic representation of the sub-tomogram average shown in N. **P-R)** Slices through electron cryo-tomograms of *∆flaA/B* cells highlighting the flagellar motor and the hook (without the filament). **S)** Sub-tomogram average of the flagellar motor and the hook structure found in *∆flaA/B* cells. **T)** Schematic representation of the sub-tomogram average shown in S. **U-W)** Slices through electron cryo-tomograms of *∆flaA/B* cells illustrating a disassembly product constituting the PL sub-complex, the rod and the hook. **X)** Sub-tomogram average of the disassembly complex shown in U-W. The dashed-orange arrow indicates the absence of the IM sub-complex in this structure. **Y)** Schematic representation of the disassembly product found in *∆flaA/B* cells. Scale bars: (red) 25 nm, (orange and white) 20 nm.

To further investigate PL sub-complexes, we generated and imaged an *S. oneidensis* strain lacking the L-ring protein FlgH. As expected, no flagella or pre-formed PL sub-complexes were seen in tomograms of *ΔflgH* cells (Movie S3). In a few cases, however, we did observe IM sub-complexes (3 examples) or the IM sub-complex with a rod and P-ring (5 examples) (Fig. 4 F-J). This indicates that the non-elaborated P-ring can form in the absence of the L-ring, but that FlgH is required for the flagellum to proceed outside the cell and for the full PL sub-complex to form. These results are consistent with studies in *E. coli* showing P-ring assembly in the absence of the L-ring^39,40^. Next, we investigated a strain lacking the flagellar filament. Previous studies showed that *S. oneidensis* cells lacking the flagellin proteins, FlaA and FlaB, are completely nonmotile^41^. In a *∆flaA/B* strain, we observed PL sub-complexes akin to those seen in wild type cells (Fig. 4 K-O). While no flagellar filaments were seen, as expected, in a few cells we observed fully-assembled motors with a hook (Fig. 4 P-T). We also frequently observed a complex comprising the PL sub-complex together with the rod and the hook, but no IM sub-complex or export apparatus, in the *ΔflaA/B* strain (examples in Fig. 4 U-Y). Again, we reasoned that this must be a disassembly intermediate, as the hook and rod proteins are secreted to the periplasm by the T3SS export apparatus. In many cases multiple copies of this disassembly product were present at the cell pole (Fig. 4W and Movie S4), suggesting an active process of attempted flagellar assembly and disassembly.

Taken together, our observations from *L. pneumophila, P. aeruginosa* and *S. oneidensis*, summarized in Figure 5, suggest an alternative model of flagellar assembly that differs from the model previously suggested for *Salmonella*^24,25^. Specifically, we found that in all three species, the P- and L-rings, together with their associated rings, formed an independent stable complex embedded in the outer membrane in the absence of other flagellar proteins. Moreover, in *L. pneumophila* and a *ΔflgH* mutant of *S. oneidensis* we found an independent IM sub-complex (C- and MS-rings with the export apparatus) embedded in the inner membrane. In *L. pneumophila*, this IM sub-complex was found in the vicinity of the PL sub-complex, suggesting a find-and-capture assembly mechanism in which the two sub-complexes form independently and come together, allowing flagellum formation to proceed. Our results suggest that once the two sub-complexes join, the transition to the fully-assembled flagellum is rapid, as reflected by the low number of particles found in intermediate states. This is in accordance with previous observations^42^.

**Figure 5:**
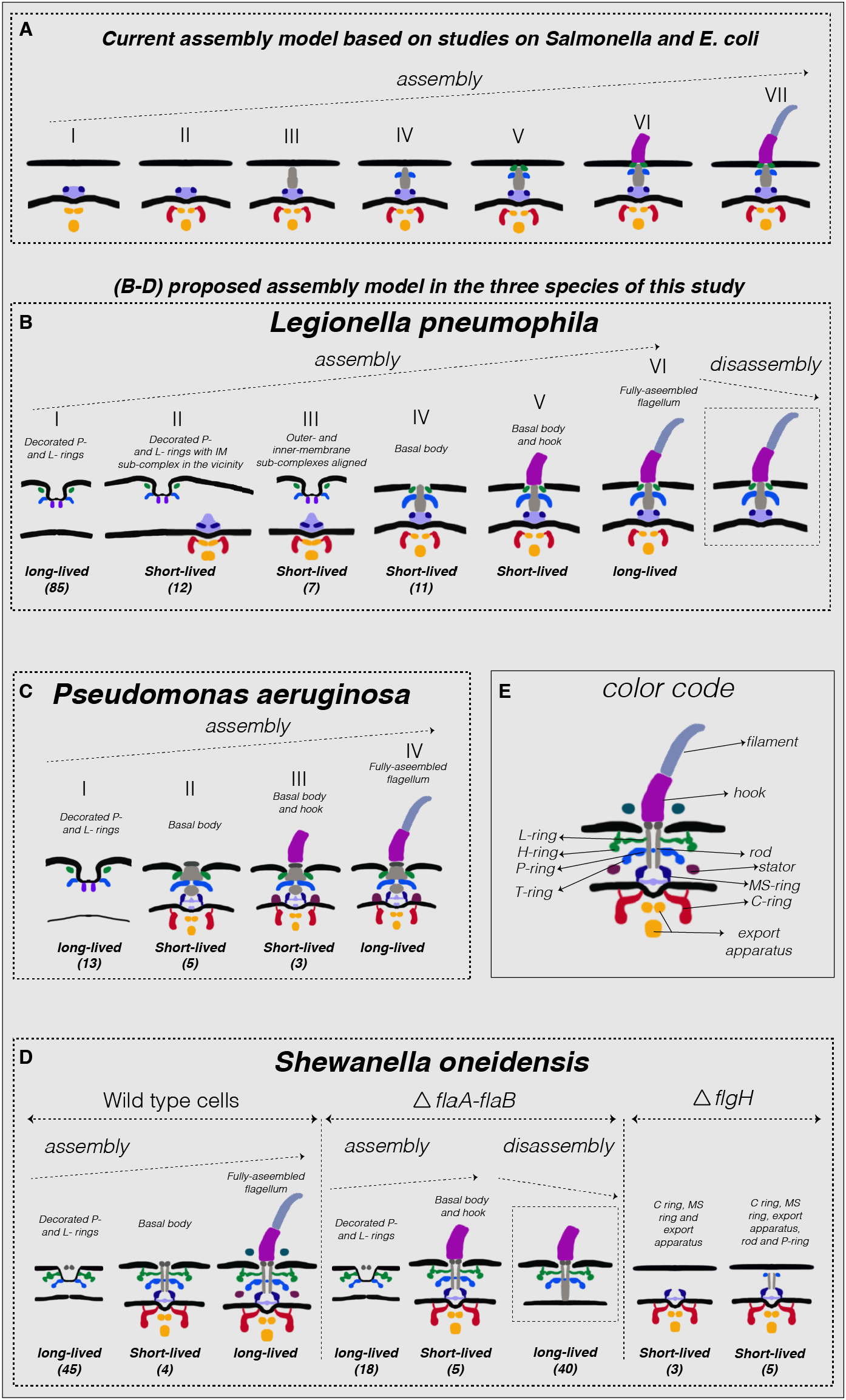
Summary of observations and proposed model of assembly. **(A)** Schematic representation of the previous model of the flagellar assembly pathway in *Salmonella* based on references^22,24–26^. **B-D)** Schematic representations of the various subcomplexes observed in this study in *L. pneumophila*, *P. aeruginosa* and *S. oneidensis* (wild type and mutant strains), respectively. Numbers in parentheses represent the number of particles observed in that particular state. In each case, the observed sub-complexes are arranged according to the assembly model proposed in the text. **E)** Labeled schematic representation of the fully-assembled flagellum in *S. oneidensis* for reference. Note that the same color code applies to all species shown.

We propose that the PL sub-complexes are the first to assemble since we observed many examples of isolated PL sub-complexes but we never observed an isolated IM sub-complex without the PL sub-complex in wild type cells. In this scenario, the IM sub-complex subsequently forms in the vicinity of the PL sub-complex and the two sub-complexes find each other, stimulating synthesis of the flagellum through the pre-formed bend in the outer membrane made by the PL sub-complex. The protein densities in the center of the PL sub-complexes extending inward might be involved in this process of locking the two sub-complexes together. Since the PL sub-complex is anchored to the peptidoglycan cell wall, we speculate that the IM sub-complex moves to find the PL sub-complex and not vice versa. This model leaves open the questions of how the two sub-complexes would assemble, and the mechanism by which they would find each other. Interestingly, this mechanism is reminiscent of the “outside-in” assembly mechanism of the closely-related T3SS injectisome in *Yersinia*, which is thought to assemble from the outside in, proceeding from the initial formation of a ring complex in the outer membrane^43^.

Previous EM studies of lysed *Salmonella* mutants revealed ring structures in the outer membrane corresponding to PL sub-complexes. These complexes were seen in mutants lacking the hook component, but not those lacking rod components^25^. It is therefore possible that the flagellar assembly mechanism of peritrichous species like *Salmonella* may differ from that of species with a single polar flagellum like the ones we imaged here.

Finally, it is possible that at least some of the PL sub-complexes we observed were the last stable complex in a disassembly process. For instance, in *L. pneumophila* cells with a large intermembrane distance, the stable PL and IM sub-complexes could have been pulled apart. For several reasons, however, we favor the idea that most of the PL sub-complexes we observed were assembly intermediates. First, in *S. oneidensis* mutants unable to fully assemble flagella, as well as in a lysed *L. pneumophila* cell, we observed clear disassembly intermediates (containing periplasmic components secreted by an export apparatus that was no longer associated with the complex). Such intermediates are consistent with those previously reported in *Caulobacter crescentus*^34–37^. The fact that we did not see any such intermediates in intact wild type cells indicates that those cells were not frequently disassembling flagella. This is unsurprising given the great energetic cost of assembling a flagellum – consuming ~2% of a cell’s total energy and ~8% of its total protein (see ^44^ and references therein) and taking a generation-time or longer to reach full length^22^. Second, we observed PL sub-complexes next to fully assembled flagella at the cell pole, and often in multiple copies (Fig. S3). Due to the energetic and temporal cost just described, we think it is unlikely that all of these cells had gone through multiple rounds of loss and assembly of their single flagellum, starting from scratch each time. In fact, a study in *Salmonella* found that cells repair flagella which are broken mechanically, only replacing them de novo if the filament is denatured using a laser pulse^45^. We think it is more likely that cells assemble multiple PL sub-complexes, perhaps to aid in the capture of the IM sub-complex.

## Acknowledgments

This work was supported by the National Institute of Health (NIH, grant R01 AI127401 to G.J.J.). M.K. is supported by a postdoctoral Rubicon fellowship from De Nederlandse Organisatie voor Wetenschappelijk Onderzoek (NWO). S.P. and M.Y.E.-N. are supported by the Air Force Office of Scientific Research Presidential Early Career Award for Scientists and Engineers (FA955014-1-0294, to M.Y.E.-N.).

## Supplementary Information

**Figure S1:**
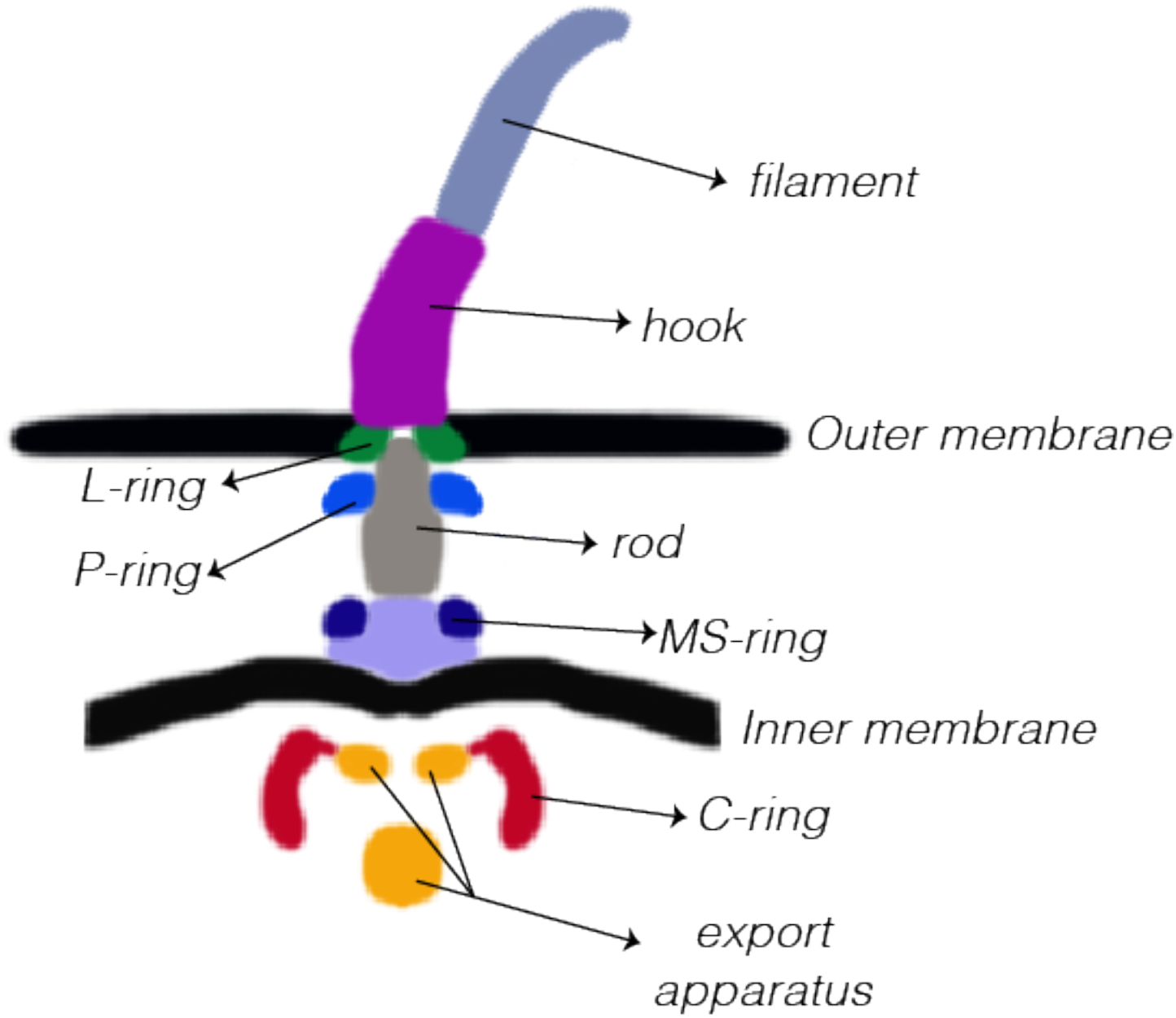
A schematic representation of the *Salmonella* flagellar motor highlighting the different parts of the structure.

**Figure S2:**
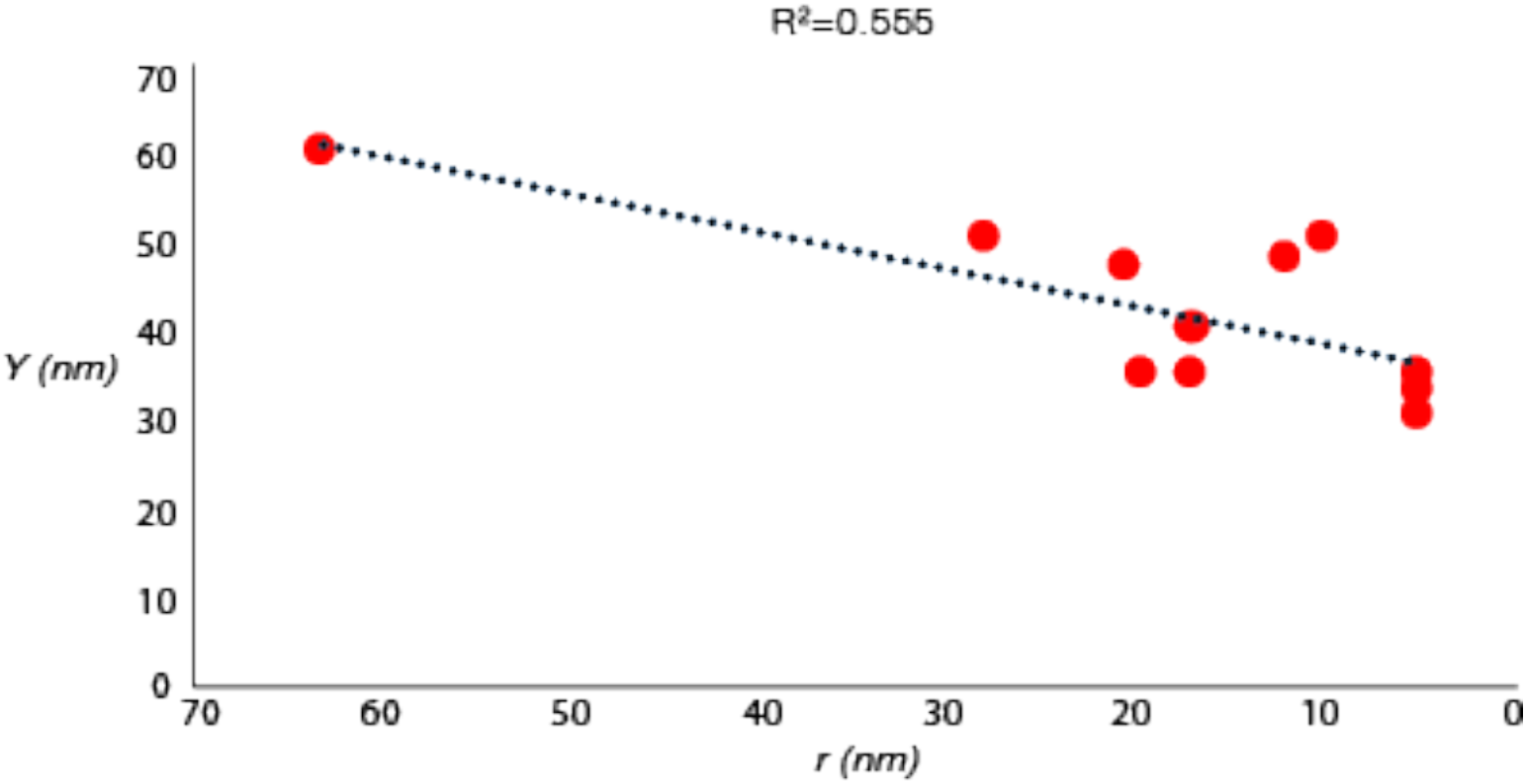
Correlation between lateral displacement of the IM and PL sub-complexes (*r*) and the distance between the inner and outer membranes (*Y*) in *L. pneumophila*. The dotted line is the trend line with the equation Y=0.4239X + 34.18.

**Figure S3:**
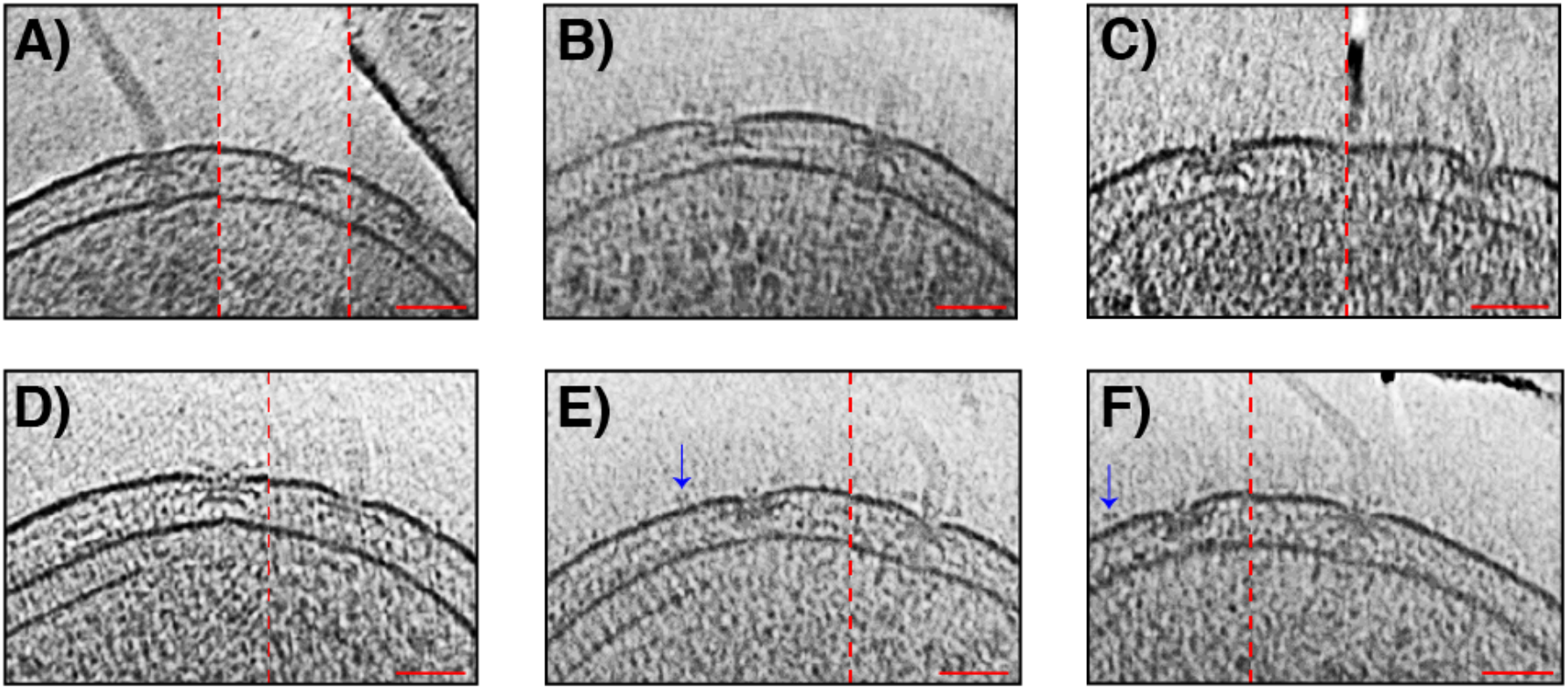
Examples of PL sub-complexes next to fully-assembled flagella in wild type *S. oneidensis* cells. Note that in some examples extracellular densities are visible next to PL sub-complexes. These extracellular densities do not appear in the sub-tomogram average as they were at different locations relative to the sub-complex and not present near all sub-complexes. They can also be seen at a distance from the PL sub-complexes (as in E and F, blue arrows) suggesting that they are not specific to PL sub-complexes. Dashed red lines indicate images that are composites of two (or more) images to show particles of interest at different z levels in the tomogram. Scale bars are 50 nm.

**Table S1:**
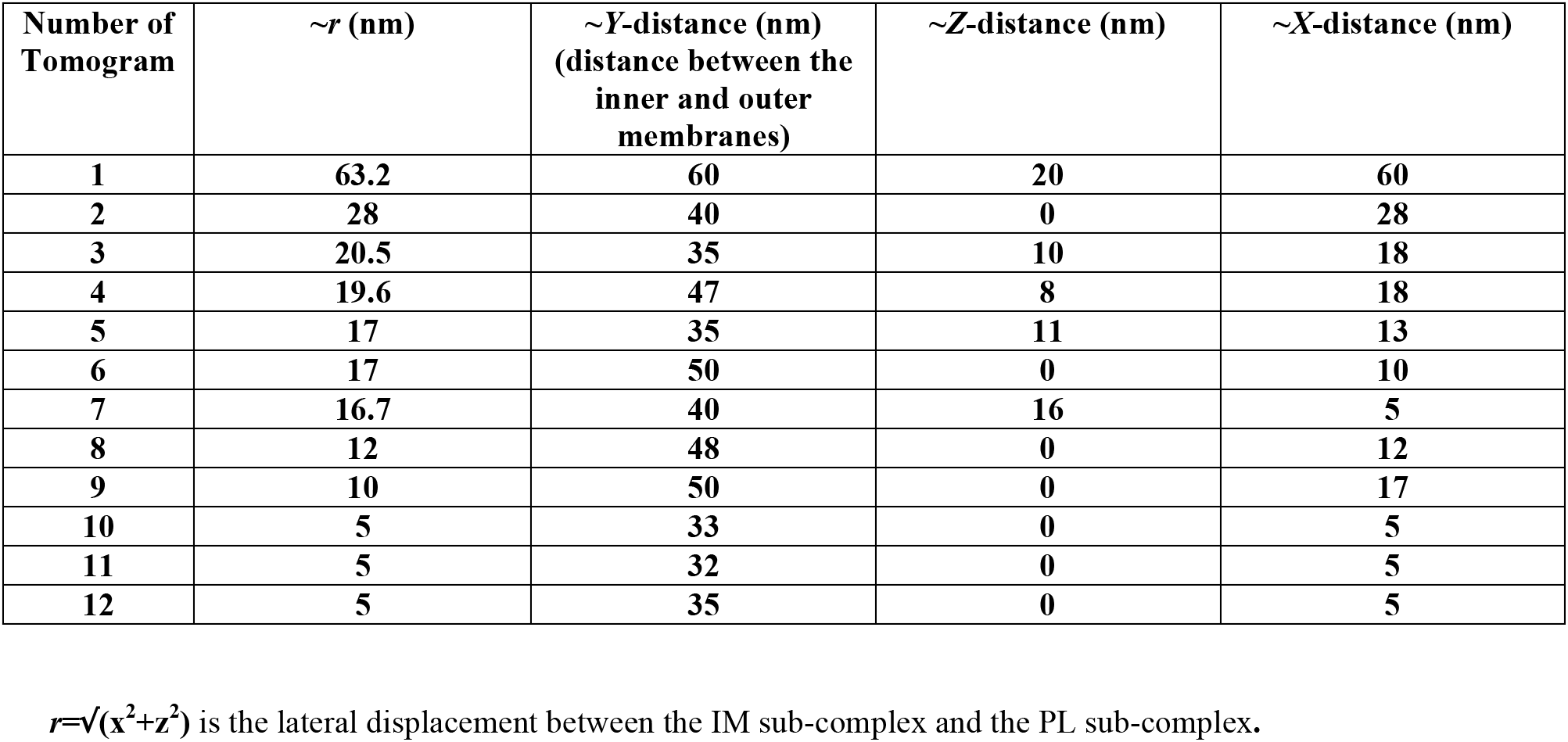
Distance between IM and PL sub-complexes in the three Cartesian axes (*x*, *y*, *z*) in *L. pneumophila*.

| Number of Tomogram | $\sim r$ (nm) | $\sim Y$-distance (nm)<br>(distance between the inner and outer membranes) | $\sim Z$-distance (nm) | $\sim X$-distance (nm) |
| --- | --- | --- | --- | --- |
| 1 | 63.2 | 60 | 20 | 60 |
| 2 | 28 | 40 | 0 | 28 |
| 3 | 20.5 | 35 | 10 | 18 |
| 4 | 19.6 | 47 | 8 | 18 |
| 5 | 17 | 35 | 11 | 13 |
| 6 | 17 | 50 | 0 | 10 |
| 7 | 16.7 | 40 | 16 | 5 |
| 8 | 12 | 48 | 0 | 12 |
| 9 | 10 | 50 | 0 | 17 |
| 10 | 5 | 33 | 0 | 5 |
| 11 | 5 | 32 | 0 | 5 |
| 12 | 5 | 35 | 0 | 5 |

$r=\sqrt{(x^2+z^2)}$ is the lateral displacement between the IM sub-complex and the PL sub-complex.

**Supplementary movies:**

**Movie S1:** Electron cryo-tomogram of a *L. pneumophila* cell highlighting the presence of independent PL and IM-sub-complexes.

**Movie S2:** Electron cryo-tomogram of a *L. pneumophila* cell highlighting the presence of independent PL and IM-sub-complexes.

**Movie S3:** Electron cryo-tomogram of a *∆flgH S. oneidensis* cell. No PL sub-complexes or flagella were seen in these cells.

**Movie S4:** Electron cryo-tomogram of a *∆flaA/flaB S. oneidensis* cell highlighting the presence of two complexes constituting the PL sub-complex, the hook and the rod.

## Material and Methods

### Strains and Growth Conditions

*S. oneidensis ∆flaA∆flaB* cells were grown using the batch culture method and *S. oneidensis* MR-1 wild-type cells were grown using either the chemostat or the batch culture methods. Detailed description of both methods can be found in^33^. Briefly, in the chemostat method, 5 mL of a stationary-phase overnight LB culture was injected into a continuous flow bioreactor containing an operating liquid volume of 1 L of a defined medium^46^, while dissolved oxygen tension (DOT) was maintained at 20%. After 20 h, and as the culture reached stationary phase, continuous flow of the defined medium^46^ was started with a dilution rate of 0.05 h^−1^ while DOT was still maintained at 20%. After 48 h of aerobic growth under continuous flow conditions, the DOT was manually reduced to 0%. O_2_ served as the sole terminal electron acceptor throughout the experiment. pH was maintained at 7.0, temperature at 30 °C, and agitation at 200 rpm. Either 24 or 40 hours after DOT reached 0%, samples were taken from the chemostat for ECT imaging.

In the batch culture method, 200 μL of an overnight LB culture of *S. oneidensis* cells was added to each of two sealed and autoclaved serum bottles containing 60 mL of a defined medium^46^. One of the two bottles acted as a control and was not used for imaging. To this control bottle, 5 μM resazurin was added to indicate the O_2_ levels in the medium. The bottles were then placed in an incubator at 30 °C, with shaking at 150 rpm until the color due to resazurin in the control bottle completely faded, indicating anaerobic conditions. At this point, samples were taken for ECT imaging from the bottle that did not contain resazurin.

The *∆flgH* mutant was constructed by a markerless in-frame deletion in the *S. oneidensis* MR-1 background made by homologous recombination using the pSMV3 suicide vector^47^ containing up- and downstream regions cloned using BamHI and SacI. The deletion was confirmed by PCR and swim-plate assay (lack of swimming on 0.3% LB agar, and complementation by plasmid-expressed FlgH) and verified by Sanger sequencing with flanking primers. Primers for the deletion construct and flanking region are as follows: HdelUpF, ACGGGATCCCGGCAACGCACAAATGATGCG, HdelUpR, CCAGTCGCTCATAAAGAACTGGCTGAGCGCAGCGGCCAATAGTAA, HdelDnF, TTACTATTGGCCGCTGCGCTCAGCCAGTTCTTTATGAGCGACTGG, HdelDnR, ACGGAGCTCGGCGCTGCACCCACTAAGTTT, HdelFlankF, GGAAGTCGTCGAAGAGGTTGGAC, HdelFlankR, CCATGCAAAGCTCCTGCCACTT. *S.oneidensis ΔflgH* cells were grown aerobically in LB culture at 30 °C to an OD_600_ of 2.4–2.8.

*L. pneumophila* Lp02 strain (*thyA hsdR rpsL*) is a derivative of the clinical isolate *L. pneumophila* Philadelphia-1. *L. pneumophila* cells were grown on ACES [*N*-(2-acetamido) −2-aminoethanesulfonic acid]-buffered charcoal yeast extract agar (CYE) or in ACES-buffered yeast extract broth (AYE). The culture media (CYE and AYE) were supplemented with ferric nitrate and cysteine hydrochloride. *L. pneumophila* Lp02 strain is a thymidine auxotroph, so cells were grown in the presence of thymidine (100 μg/ml). Cells were grown to early stationary phase (OD_600_ ~2.5) and subsequently harvested for ECT sample preparation.

*Pseudomonas aeruginosa* PAO1 cells were grown on LB plates overnight at 37 °C. After that, cells were inoculated into 5 ml MOPS [(3-(*N*-morpholino) propanesulfonic acid)] Minimal Media Limited Nitrogen and grown for ~24 hours at 30 °C.

### ECT sample preparation and imaging

BSA-treated 10-nm colloidal gold solution was mixed with cells from the three species and 4 μL of this mixture was applied to a glow-discharged, carbon-coated, R2/2, 200 mesh copper Quantifoil grid (Quantifoil Micro Tools) in a Vitrobot chamber (FEI). Excess liquid was blotted off and the grid was plunge frozen for ECT imaging. Imaging of all ECT samples of *S. oneidensis* and *P. aeruginosa* was performed on an FEI Polara 300-keV field emission gun electron microscope (FEI company, Hillsboro, OR, USA) equipped with a Gatan image filter and K2 Summit direct electron detector in counting mode (Gatan, Pleasanton, CA, USA). *L. pneumophila* cells were imaged using an FEI Titan Krios 300 kV field emission gun transmission electron microscope equipped with a Gatan imaging filter and a K2 Summit direct detector in counting mode (Gatan). Data were collected using the UCSF Tomography software^48^, with each tilt series ranging from −60° to 60° in 1° increments, and an underfocus of ^~^5–10 μm. A cumulative electron dose of ^~^130–160 e^-^/A^2^ for each individual tilt series was used for *S. oneidensis* and *P. aeruginosa* while a cumulative dose of *~* 100 e^-^/A^2^ was used for *L. pneumophila*.

### Image processing and sub-tomogram averaging

The IMOD software package was used to calculate three-dimensional reconstructions of tilt series^49^. Alternatively, the images were aligned and contrast transfer function corrected using the IMOD software package before producing SIRT reconstructions using the TOMO3D program^50^. Sub-tomogram averages with 2-fold symmetrization along the particle Y-axis were produced using the PEET program^51^.

